# A Framework for Cortical Layer Composition Analysis using Low Resolution T1 MRI Images (August 2018)

**DOI:** 10.1101/390112

**Authors:** Ittai Shamir, Omri Tomer, Zvi Baratz, Galia Tsarfaty, Maya Faraggi, Assaf Horowitz, Yaniv Assaf

## Abstract

The layer composition of the cerebral cortex represents a unique anatomical fingerprint of brain development, function, connectivity and pathology. Historically the cortical layers were investigated solely ex-vivo using histological means, but recent magnetic resonance imaging (MRI) studies suggest that T1 relaxation images can be utilized to separate the layers. Despite technological advancements in the field of high resolution MRI, accurate estimation of whole brain layer composition has remained limited due to partial volume effects, leaving some layers far beyond the image resolution. In this study we offer a simple and accurate method for layer composition analysis, resolving partial volume effects and cortical curvature heterogeneity. We use a low resolution echo planar imaging inversion recovery (EPI IR) MRI scan protocol that provides fast acquisition (~12 minutes) and enables extraction of multiple T1 relaxation time components per voxel, which are assigned to types of brain tissue and utilized to extract the subvoxel composition of each T1 layer. While previous investigation of the layers required the estimation of cortical normals or smoothing of layer widths (similar to VBM), here we developed a sphere-based approach to explore the inner mesoscale architecture of the cortex. Our novel algorithm conducts spatial analysis using volumetric sampling of a system of virtual spheres dispersed throughout the entire cortical space. The methodology offers a robust and powerful framework for quantification and visualization of the layers on the cortical surface, providing a basis for quantitative investigation of their role in cognition, physiology and pathology.

## I. Introduction

Study of the laminar structure of the cerebral cortex was first made possible in the beginning of the 20th century ex-vivo through the use of histological methods [1]. Years later, in 1950, renowned neuroanatomist Gerhardt von Bonin famously stated in an article on the cerebral cortex that “the cortex is both chaos and order, and therein lies its strength” [2]. On the chaos side, the cerebral cortex has a highly tortuous surface consisting of many gyri and sulci with an overall thickness that varies regionally between 2 and 4 mm on average throughout cortical regions. On the order side, the cortex is characterized by a highly organized laminar structure consisting of six cortical layers, each characterized by different types of neurons. While the order of the cortical layers remains constant, the thickness of each layer varies regionally throughout cortical regions and therein lies its “strength”. The layers are initially formed during brain development, when neurons migrate to form the cortex, playing an important role in brain connectivity and function. The layers and their composition are assumed to play an integral role in the function, development and pathology of the brain [3].

With the advent of MRI, visualization of the overall cortical thickness using T1-weighted MRI images has been successfully achieved. Cortical thickness visualization includes accurate delineation of the inner cortical surface, bordering with myelin rich white matter, and the outer cortical surface, bordering with pia matter and the surrounding cerebral spinal fluid [4], [5]. Recent studies suggest that different MRI imaging protocols can also be utilized to provide layer-specific information, using either high resolution at high magnetic field or subvoxel modelling at lower resolutions. These approaches use a variety of MRI images, including T1, T2 and T2^∗^ weighted images, as well as R1, R2 and R2^∗^ susceptibility images [6], [7], [8], [9], [10], [11], [12].

The T1 MRI approach to imaging the substructure of the cortex has shown great potential. An early IR MRI study of both rat and human cortices revealed six T1 clusters, corresponding to six cortical layers [15]. A larger scale follow-up study of human cortices using the same IR MRI protocol revealed that the six clustered T1 components along the cortex are similar throughout subjects [14]. Still, automation of the visualization process of the cortical layers remains one of the most significant neuroimaging challenges of recent years.

Visualization of the cortical layer components remains hindered by two main imaging challenges. The first challenge involves partial volume effect (PVE), an imaging effect occurring when voxel size exceeds the size of tissue detail [13]. Our premise is that the solution to imaging the cortical layers does not necessarily lie in increasing T1 MRI resolution, since even such high resolution images are afflicted by PVE, posing a limiting factor in visualization of the layers [14]. The second challenge involves the intricate geometry of the cortex, which has been typically approached either by estimating normals to the cortical surfaces and investigating the layer composition along them or by smoothing layer widths, similar to voxel-based morphometry (VBM) [15], [27], [28]. The applicability of such approaches is limited, not only because minor errors in cortical surface estimation can lead to greater errors in normal estimations, but also because this process has to be repeated accurately throughout the entire cortex.

In this work we present a method for investigating the cortical layers through subvoxel modelling at lower resolution T1 MRI. We use a low resolution echo planar imaging inversion recovery (EPI IR) protocol that provides fast acquisition (~12 minutes) and enables extraction of multiple T1 components per voxel, which are assigned to brain tissue types and utilized to extract the subvoxel composition of each T1 layer [14], [15], [16], [17]. We then explore the mesoscale laminar architecture of the cortex using a sphere-based approach, implementing a geometric solution based on cortical volume sampling using a system of virtual spheres dispersed throughout the entire cortex. A spherical shape was chosen due to its symmetry and invariance to rotation, offering a simple and robust alternative to cortical normals.

Our methodology offers an automated and unbiased whole-brain solution to investigating the complex mesoscale laminar architecture of the cortex. We suggest that our powerful tool for investigating the layers could enable expansion of studies on the role of cortical thickness in brain function and behavior to the cortical layer level.

## II. Methods and Materials

### A. Subjects

Fifteen healthy human subjects were recruited for this study (N=15), including 7 male and 8 female, 23-43 year old, all right handed. Subjects were neurologically and radiologically healthy, with no history of neurological diseases, and normal appearance of clinical MRI protocol. The imaging protocol was approved by the institutional review boards of Sheba Medical Centers and Tel Aviv University, where the MRI investigations were performed. All subjects provided signed informed consent before enrollment in the study.

### B. MRI Acquisition

All experiments were scanned on a 3T Magnetom Siemens Prisma (Siemens, Erlangen, Germany) scanner with a 64- channel RF coil.

Two T1 weighted MRI sequences were used in order to characterize the cortical layers:

1. *An inversion recovery echo planar imaging (IR EPI) sequence*, with the following parameters: TR/TE=10000/30 ms and 60 inversion times spread between 50 ms up to 3,000 ms, voxel size 3×3×3 *mm*^3^, image size 68×68×42 voxels, each voxel fitted with up to 7 T1 values [14] (see below). The acquisition time for the inversion recovery data set was approximately 12 min.
2. *An MPRAGE sequence*, with the following parameters: TR/TE=1750/2.6 ms, TI=900 ms, voxel size 1×1×1 *mm*^3^, image size 224×224×160 voxels, each voxel fitted with a single T1 value. This sequence was used as an anatomical reference with high gray/white matter contrast.

### C. IR Decay Function Fit

These data were used for multiple T1 analysis, by calculating T1 values and their corresponding partial volumes on a voxel-by-voxel basis. The IR datasets were fitted to the conventional inversion recovery decay function with up to 7 possible T1 components per voxel [14]:

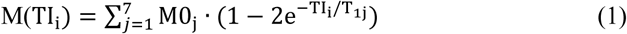

Where:

M(TI_i_)- Magnetization at the *i*^th^ inversion recovery image, in other words the magnetization measured for each specific T1 component.

M0_j_- Predicted magnetization at TI=0ms for each T1 component (j) in the voxel.

T_1j_- Longitudinal relaxation time for each T1 component.

j was set up to 7 for the low resolution experiments, indicating fit to seven individual exponential fits, based on the assumption that there are 7 T1 components in the tissue – 1 for CSF, 1 for WM and heavily myelinated layer of the cortex and additional 5 cortical layers.

Normalization of each of the predicted magnetization values according to 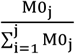 then represents the voxel contribution of each corresponding T1 component (j).

### D. T1 Probabilistic Classification

In order to accurately classify whole brain T1 values to their corresponding brain tissues, a T1 histogram was initially plotted. Each IR EPI set of images consists of up to ~1.36×10^6^ potential T1 values: 68×68×42 [voxels] ×7 [T1 fitted values]. Each voxel was given a weight greater than one in order to accurately represent all its fitted T1 values according to their corresponding partial volume contributions. The T1 histogram was then fitted to a probabilistic mixture model (similarly to the method shown in [14], [15]), consisting of t-distributions [16], [17]. The probability of each t-distribution in the voxel was calculated using Bayes’ formula:

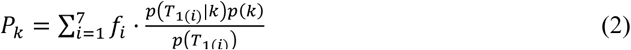

Where:

*k*- A specific t-distribution.

*T*_1_(*i*)- T1-value of the *i*^th^ component of the voxel.

*f*_*i*_- Partial volume of *T*_1_(*i*) (normalized as show in previous section).

*p*(*T*_1_)- General whole-brain probability of a *T*_1_-value.

*p*(*k*)- Probability of t-distribution k.

*p*(*T*_1_|*k*)- Probability of the *T*_1_-value in t-distribution *k*.

This model was used as a means of probabilistic classification of T1 values per subject into 11 clusters, corresponding to different types of brain tissue (see figure 1):

**Fig. 1.**
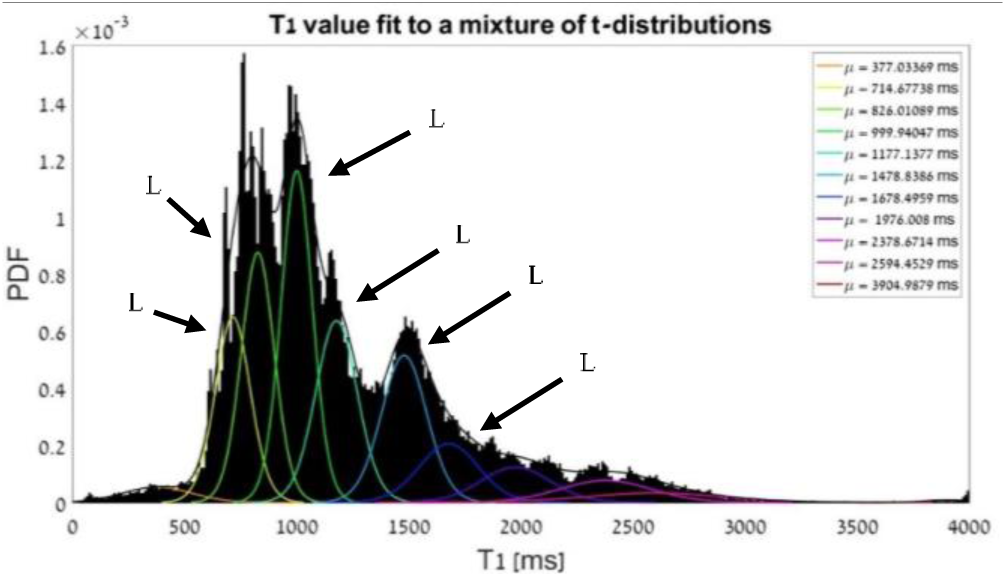
T1 histogram (black) fit to a mixture of 11 t-distributions, where μ_*k*_ represents the expected value of each distribution, or the center of its curve. Overall fit is represented by the black outline, individual t-distributions represented according to color map, where t-distributions 2,3,4,5,6 and 7 correspond to T1 layers 6,5,4,3,2 and 1 (respectively).

1. *White matter (WM)* characterized by low T1 values, represented by 1^st^ t-distribution.
2. *Grey matter (GM)* characterized by mid-range T1 values, represented by 2^nd^-7^th^ t-distributions corresponding to 6 T1 layers, with increasing degrees of myelination.
3. *Cerebral spinal fluid (CSF)* characterized by high T1 values, represented by 8^th^-11^th^ t-distributions.

### E. Image Registration

In the interest of accurately sampling T1 layer probability maps inside the cortical spheres, all T1 layer probability maps were registered to the anatomical MPRAGE image.

Registration was completed in SPM12 using a rigid body transformation with the first IR image (IR1) as the source image.

### F. Cortical Volume Sampling

In order to provide accurate volumetric sampling of the complex geometry of the cortex, we offer a simple geometric alternative to cortical normals-creating a sampling system of virtual spheres filling the entire cortex volume. A spherical shape was chosen as due to its symmetry and invariance to rotation, minimizing errors in cortical normals associated with surface calculations and consequently enabling more accurate sampling and localization throughout the cortex. The following steps were taken to create and implement this volumetric sampling system:

1. Cortical surface delineation: The anatomical MPRAGE image was segmented in the *FreeSurfer* pipeline [18], revealing the following three cortical surfaces:
  a. Inner surface-cortical GM bordering with the underlying WM.
  b. Mid surface-an estimation of the center of cortical GM, based on the inner and outer surfaces.
  c. Outer surface-cortical GM bordering with the surrounding CSF.
2. Volumetric sampling system: Each cortical surface is represented by a triangular mesh, consisting of vertices corresponding to points in space (~150,000 per hemisphere) and faces corresponding to three vertices per triangle. Each sphere was centered on a vertex on the mid surface, with its top tangential to the outer surface and its bottom tangential to the inner surface. Since the average sphere radius is ~1mm and the average distance between adjacent vertices on the surface is ~1mm, volumetric overlap between spheres occurs. In order to sample the entire cortex and yet avoid excessive volumetric overlap, sampling rate was chosen by subsampling the mid surface vertices by a factor of 2, to ~75,000 spheres per hemisphere (see figure 2).
3. Data sampling implementation: One of the main challenges in implementing spherical sampling of low resolution data lies in the resolution difference between the spherical sampling system (average radius of ~1 mm) and the T1 data ( 3^3^ *mm*^3^). Overcoming this resolution gap demanded a super-resolution solution based on estimation of subvoxel information. Our offered solution includes the following steps:
  a. *Partitioning each voxel into subvoxels*: Degree of partitioning was chosen so that each subvoxel has the largest volume while still enabling accurate group representation of a spherical volume with a radius of 1 mm. Consequently, each voxel was partitioned into 10^3^ subvoxels, each assigned location properties, primarily its location inside or outside of a given sphere.
  b. *Assignment of spherical volume weights*: Each sphere in our sampling system was assigned weights corresponding to each voxel’s volume contribution to its spherical volume (see figure 3 below), according to the following equation:

**Fig. 2.**
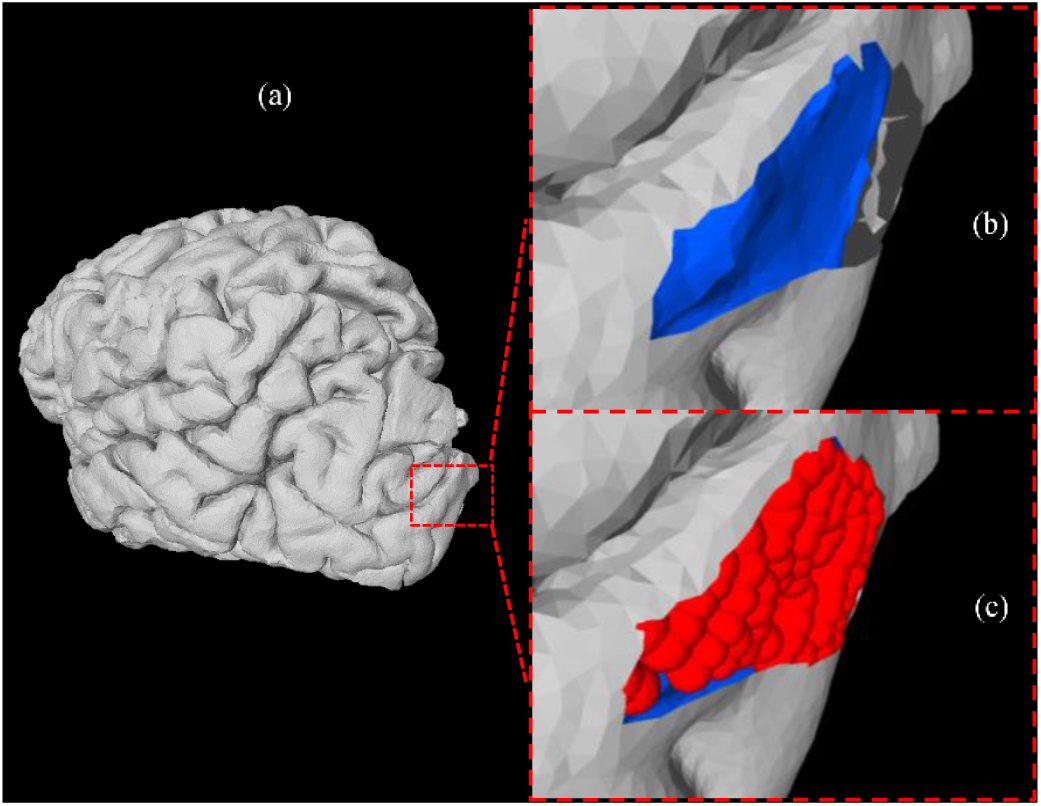
Cortical volume sampling: left side view of left hemisphere, where the outer cortical surface is represented in grey (a), the underlying inner cortical surface is represented in blue (b) and the cortical volume sampled between the two is represented by red spheres (c).

**Fig. 3.**
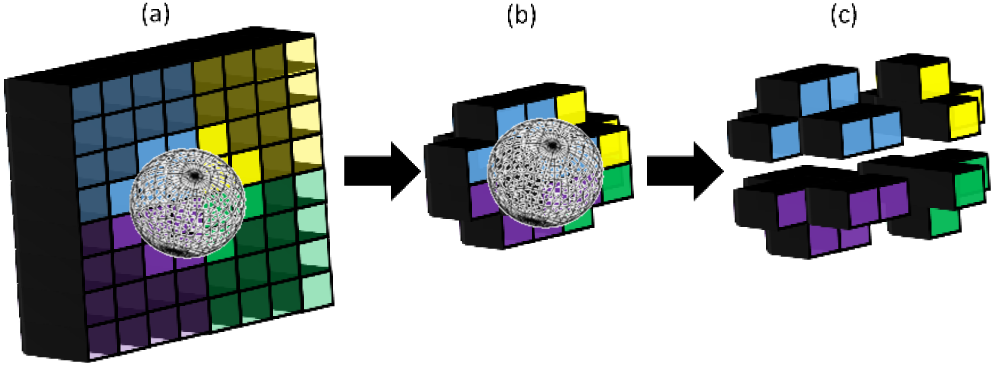
Schematic representation of voxel partitioning. A sphere is located between 4 voxels, shown in blue, yellow, green and purple (a). Each voxel is divided into 4^3^ subvoxels, leaving only those contributing to the volume of the sphere (b). The sphere’s weights are assigned according to each voxel’s contributing portion of subvoxels to its volume (c).

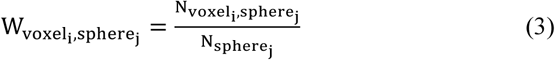 Where: W_voxel_i_,sphere_j__- Volume weight of voxel i per sphere j. N_voxel_i_,sphere_j__- Number of subvoxels from voxel i located inside sphere j. N_sphere_j__- Total number of subvoxels located inside sphere j.
  c. *Cortical composition analysis inside sphere*: Volume weights were multiplied by their corresponding voxel probability maps (see *T1 probabilistic classification*) per each sphere throughout our entire sampling system, according to the following equation:

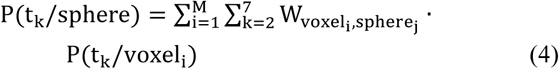 Where: P(t_k_/sphere)- Probability of t-distribution k per sphere. k-t-distributions 2,3,..,7, representing T1 layers 6,5,..,1 (respectively). M-Number of voxels within which sphere j lies. W_voxel_i_,sphere_j__ - Volume weight of voxel i per sphere j. P(t_k_/voxel_i_)- Probability of t-distribution k in voxel i.

## III. Results

Use of the cortical spheres enabled a rotationally invariant estimation of layer composition within the cortex with simple projection of the quantitative layer width onto the surface. In other words, implementation of our methodology resulted in 6 multidimensional datasets per subject, each consisting of ~150,000 spheres dispersed throughout both hemispheres and revealing individual T1 layer compositions.

The results were visualized by projecting each T1 layer composition onto the subject’s cortical mid surface. In order to better conduct exploratory spatial analysis of each layer composition, intra-subject values were averaged regionally throughout 75 different cortical regions, using *FreeSurfer*’s automatic surface-based parcellation atlas Destrieux [19] (see figure 4).

**Fig. 4.**
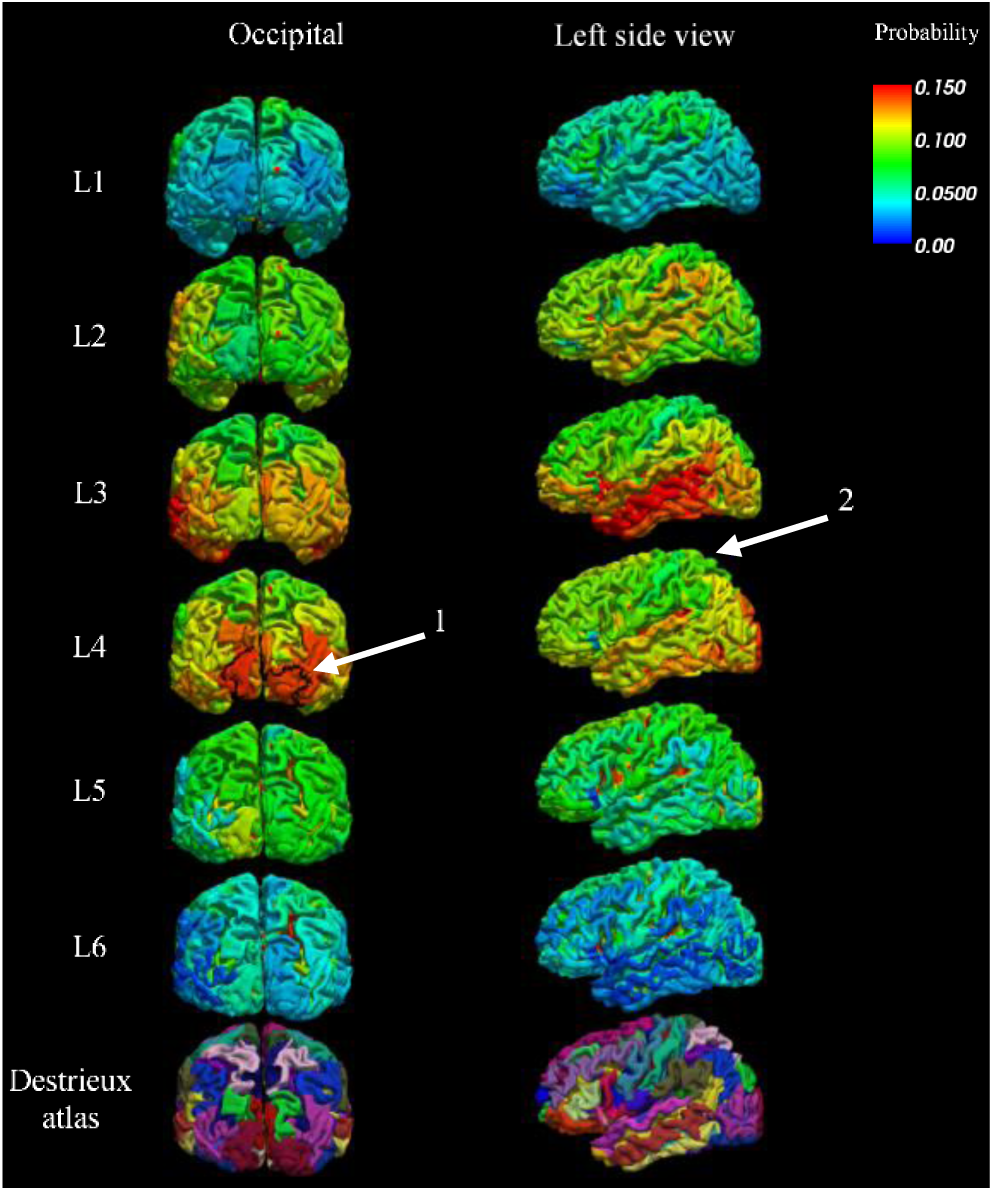
Single subject T1 layer probability maps (L1-L6), averaged per each of the 75 regions in Destrieux Atlas. Arrows indicate interesting features: arrow 1 indicates high T1 layer 4 presence in the primary visual area (V1 outlined in black), and arrow 2 indicates lower presence in the motor cortex.

The process was repeated for 15 subjects and then inter-subject cortical composition values were averaged again per each of the 75 Destrieux atlas regions. Results were visualized by projecting each averaged layer composition onto a cortical surface representing the average brain of all 15 subjects (see figure 5 below).

**Fig. 5.**
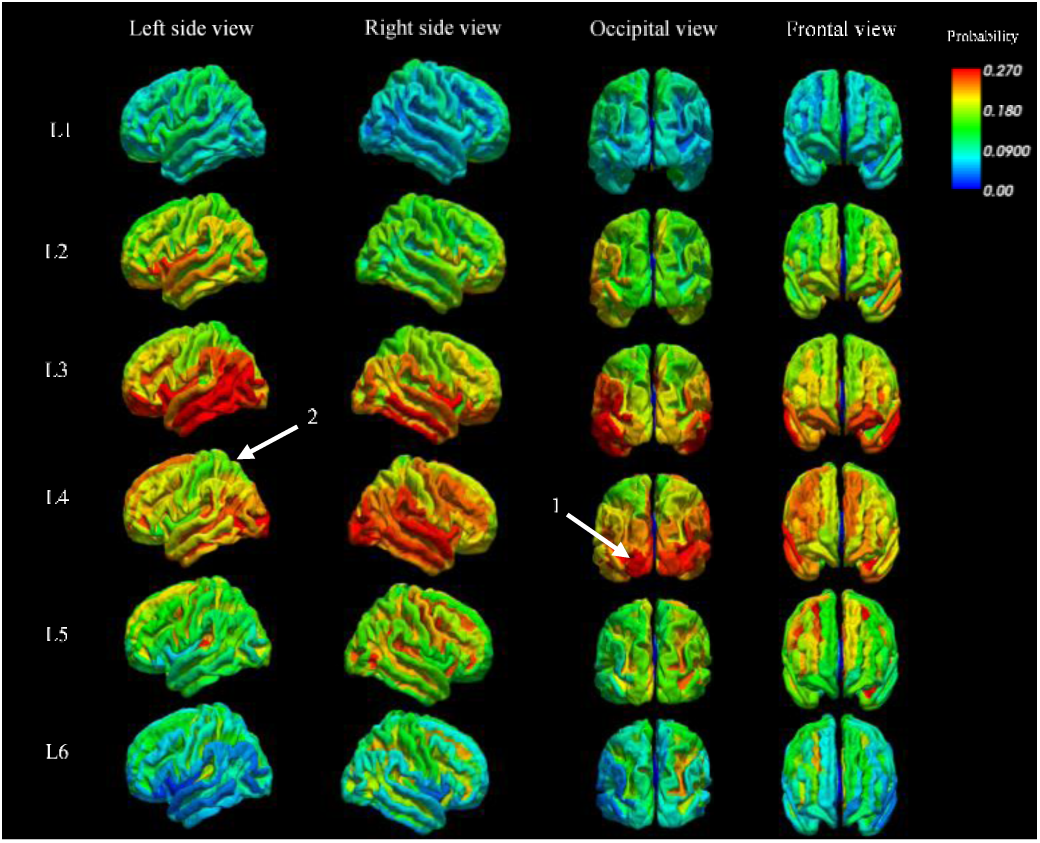
15 subject average T1 layer probability maps (L1-L6), averaged per each of the 75 regions in Destrieux Atlas. Arrows indicate the recurring interesting features from single subject analysis (see figure 4). Arrow 1 indicates high T1 layer 4 presence in the primary visual area and arrow 2 indicates lower presence in the motor cortex.

Visual assessment of these surface projections reveals that the innermost and the outermost layers, corresponding to T1 layers 1 and 6, exhibit the lowest compositional values in comparison with T1 layers 2, 3, 4 and 5. More specifically, an interesting feature recurring across subjects is the high intensity of T1 layer 4 in the primary visual area (V1), accompanied by low intensity in the motor cortex.

Quantitative assessment of the T1 layer compositions was conducted by registering the 15 subject average brain to a granularity atlas and investigating the T1 layer composition across indices. This atlas divides the cortex into regions with 6 increasing granularity indices, from the least granular allocortex regions up to the granular cortical regions (segmentation was completed based on a similar process shown in [20]). T1 layer compositions were averaged per granular index (see figure 6 below).

**Fig. 6.**
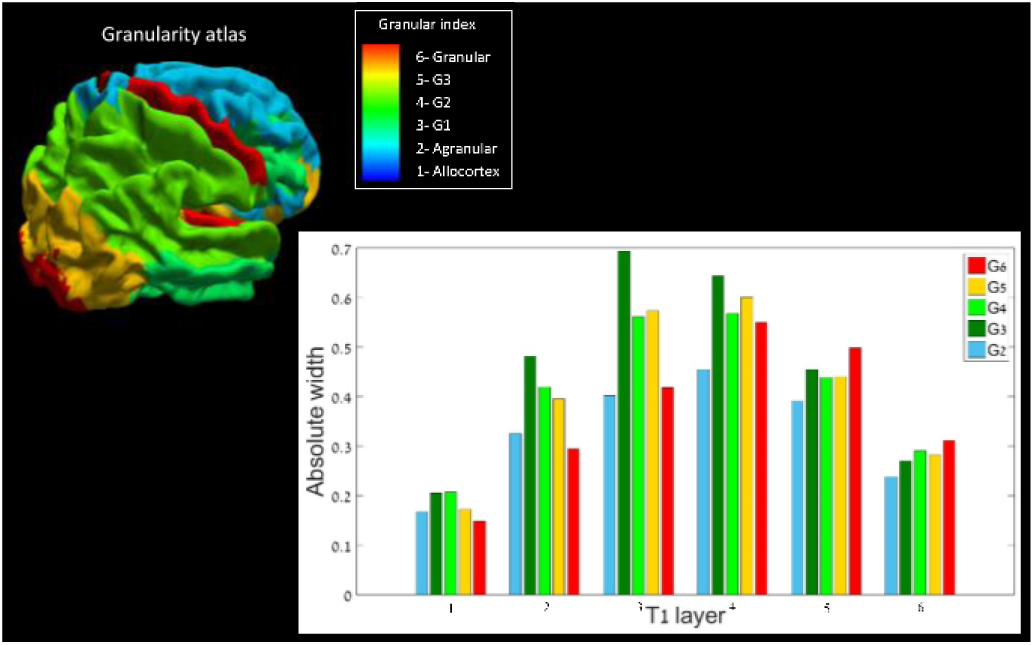
Distribution of granularity indices (G2-G6) across T1 layers throughout the cerebral cortex (right). Results are based on 15 subject average T1 layer probability maps (see figure 5), registered to a granularity index atlas (left). Allocrtex (G1) was not included in analysis, not only because it characteristically consists of fewer than 6 cortical layers, but also because of its relatively minor contribution to the cortical volume.

Once again, T1 layers 2-5 exhibit a more dominant presence throughout the cortex, compared to layers 1 and 6. T1 layers which are closer to white matter both physically as well as in cytoarchitecture (T1 layers 3-6), exhibit an increasingly high presence in more granular regions (G3-G6), while the outermost less myelinated T1 layers (layers 1 and 2) exhibit a decreasing presence in such granular regions.

## IV. Discussion

In this work we propose an automated and unbiased methodology for whole-brain investigation of the complex mesoscale laminar architecture of the cortex. The methodology is based on a surface-wise T1 layer composition analysis of inversion recovery echo planar imaging (IR EPI), which enables extraction of multiple T1 relaxation time components per voxel. The novelty lies in the use of rotationally invariant cortical spheres, sampling the regionally varying cortical thickness across the tortuous cortical folding. The selectivity of T1 measures to the cortex combined with conventional cortical surface estimation, on which our sampling system of spheres is based, overcome the low resolution of the acquired IR MRI and allow representation of layer widths in a 3D in objective manner.

While MRI visualization of the overall cortical thickness has been successfully achieved [4, 5], to this day there lacks a single automatic whole-brain approach for investigating the laminar structure of the cortex. There are two main opposing approaches for investigating the cortex through the use of T1 MRI. The first approach uses high resolution at high magnetic field, typically at the level of laminar width but on a small volume of the brain [7], [8], [14], [21], [22], [23], [24], [25], [26]. The second approach uses whole-brain subvoxel modelling at lower resolutions, which creates a localization challenge [14], [15]. The reason for the limited applicability of the abovementioned approaches is twofold. On one hand is PVE, when cortical layer detail exceeds even the resolution of high resolution images of the cortex [13], [14]. PVE is commonly dealt with by smoothing layer widths (similar to VBM), which decreases level of tissue detail [15]. On the other hand is the localization challenge inside the complex geometry of cortex, which demanded the estimation of cortical normals [15, 27, 28], a semi subjective process prone to errors due to inaccuracies in surface modelling [14].

Here we achieve super-resolution by estimating subvoxel information from a multi-T1 dataset inside a sampled system of volumetric cortical spheres. Our method has several main advantages: 1) Use of a standard MRI protocol and scanner setup (IR EPI and MPRAGE); 2) Quick acquisition of a low resolution dataset (~12 minutes); 3) Investigation of the laminar structure of the cortex in an automated whole-brain 3D manner, overcoming both PVE as well as the need for cortical normals; 4) Enabling easy visualization of laminar composition through the use of laminar surface projections; 5) Demonstration of transition between various brain regions solely based on laminar composition. Most notably, our method demonstrates accurate delineation of the primary visual cortex (V1) (see figures 4 and 5). V1 is considered a hallmark of unique cortical lamination in the visual system, which has been previously delineated successfully through the use of high resolution region-specific T1 MRI, focusing on the stripe of Gennari [8], [21], [23]. In other words, despite the low resolution of our T1 dataset, whole-brain surface projections of the layer probability maps demonstrate delineation or brain regions that have previously been delineated by high resolution investigation.

It is important to note that when discussing our results we use the term ‘T1 layer’, avoiding the traditional term ‘cortical layer’. The reasoning behind this phrasing is that T1 is not considered a direct measure of cytoarchitecture. Although multi-T1 methods investigate the laminar structure of the cortex, even high resolution T1 MRI averages contributions of many cells, including but not limited to neurons and glial cells [8], [15].

The main limiting factor of our method is the number of TIs chosen in the IR EPI sequence, which has significant impact on the probabilistic classification to T1 layers. Although our method is surface-based, it less affected by the accuracy of cortical surface modelling, thanks to the spheres’ invariance to rotation. Nonetheless, the surface-based nature of our approach can still be considered a minor limiting factor. These limiting factors can easily be dealt with by increasing the number of TIs and thus potentially increasing the accuracy of T1 layer classification, or by increasing the resolution of the anatomic MPRAGE image (from 1mm to 600um, for example), thus increasing the accuracy of the cortical surfaces. Choice of these image acquisition parameters is an act of checks and balances, with acquisition time and image resolution on either sides of the scale.

The automatic whole-brain applicability of the method could enable future analysis of large populations. Such larger scale studies of groups of healthy subjects could provide a closer look at both the statistics of T1 layer characteristics, as well as regional layer transition effects, including compositional layer variability between the gyral wall and the cap of the gyri.

It has long been accepted that the laminar structure of the cortex plays an integral role in cognition, physiology and different pathologies. For instance, certain forms of epilepsy have been linked to cortical dysplasia, a pathology involving abnormalities of the laminar structure of the cortex, which could be further studied using in-vivo subcortical imaging [29]. We suggest that use of the robust automatic tool presented here could potentially be used as a framework for quantitative expansion of such studies into the role of cortical thickness in brain function and behavior to the cortical layer level.

## References

[1] K. Brodmann, and L. J. Garey, “Brodmann’s localization in the cerebral cortex”, Springer, New York, NY, 2006 pp. 1–58.

[2] G. Von Bonin, and C. C. Thomas, “Essay on the cerebral cortex”, Springfield, ed. 3, 1950.

[3] J. Kiernan and N. Rajakumar, “Barr’s The Human Nervous System – An Anatomical Viewpoint”, Lippincott Williams & Wilkins, vol. 13, no. 62, pp. 213–246, 2014.

[4] B. Fischl and S. M. Dale, “Measuring the Thickness of the Human Cerebral Cortex from Magnetic Resonance Images”, PNAS, vol. 97, no. 20, pp. 11050–11055, 2010.

[5] L. H. Scholtens, M. A. de Reus, and M. P. van den Heuvel, “Linking Contemporary High Resolution Magnetic Resonance Imaging to the Von Economo Legacy: A Study on the Comparison of MRI Cortical Thickness and Histological Measurements of Cortical Structure”, Human Brain Mapping, vol. 36, pp. 3038–3046, 2015.

[6] R. Shafee, R. L. Buckner, and B. Fischl, “Gray matter myelination of 1555 human brains using partial volume corrected MRI images”, NeuroImage, vol. 105, pp. 473–485, 2015.

[7] J. H. Duyn, P. van Gelderen, T. Q. Li, J. A. de Zwart, A. P. Koretsky and M. Fukunaga, “High-field MRI of brain cortical substructure based on signal phase”, PNAS, vol. 104, no. 28, pp. 11796–11801, 2007.

[8] E. L. Barbier, S. Marrett, A. Danek, A. Vortmeyer, P. van Gelderen, J. Duyn, P. Bandettini, J. Grafman, and A. P. Koretsky, “Imaging Cortical Anatomy by High-Resolution MR at 3.0T: Detection of the Stripe of Gennari in Visual Area 17”, Magnetic Resonance in Medicine, vol. 48, pp. 735–738, 2002.

[9] H. Bridge, and S. Clare, “High-resolution MRI: in vivo histology?”, Philosophical Transactions of the Royal Society, vol. 361, pp.137–146, 2006.

[10] M. F. Glasser, M. S. Goyal, T. M. Preuss, M. E. Raichle, and D. C. Van Essen, “Trends and properties of human cerebral cortex: Correlations with cortical myelin content”, NeuroImage, vol. 93, pp. 165–175, 2014.

[11] A. Deistung, A. Schafer, F. Schweser, U. Biedermann, R. Turner, and J. R. Reichenbach, “Toward in vivo histology: A comparison of quantitative susceptibility mapping (QSM) with magnitude-, phase-, and R2*-imaging at ultra-high magnetic field strength”, NeuroImage, vol. 65, pp. 299–314, 2013.

[12] A. Lutti, F. Dick, M. I. Sereno, and N. Weiskopf, “Using high-resolution quantitative mapping of R1 as an index of cortical myelination”, NeuroImage, vol. 93, pp. 176–188, 2014.

[13] M. A. Gonzalez Ballester, A. P. Zisserman, and M. Brady, “Estimation of the partial volume effect in MRI”, Medical Image Analysis, vol. 6, pp. 389–405, 2002.

[14] S. Lifshits, O. Tomer, I. Shamir, D. Barazany, G. Tsarfaty, S. Rosset, and Y. Assaf, “Resolution considerations in imaging of the cortical layers”, Neuroimage, vol. 164, pp. 112–120, 2018.

[15] D. Barazany, and Y. Assaf, “Visualization of Cortical Lamination Patterns with Magnetic Resonance Imaging”, Cerebral Cortex, vol. 22, pp. 2016–2023, 2012.

[16] D. Peel, and G. J. McLachlan, “Robust mixture modelling using the t distribution”, Statistics and computing, vol. 10(4), pp. 339–348, 2000.

[17] O. Tomer, Z. Baratz, I. Shamir, D. Kaptzon, A. Horowitz, M. Faraggi, D. Barazany, and Y. Assaf, “A probabilistic method for modelling cortical layer composition in sub-voxel resolution”, Poster session presented at the Organization for Human Brain Mapping Association, Singapore, June, 2018. Available: https://files.aievolution.com/hbm1801/abstracts/32677/2510_Tomer.pdf

[18] B. Fischl, “FreeSurfer”, NeuroImage, vol. 62, pp. 774–781, 2012.

[19] B. Fischl, and A. van der Kouwe, “Automatically Parcellating the Human Cerebral Cortex”, Cerebral Cortex, 14, 11-22, 2004.

[20] S. F. Beul, and C. C. Hilgetag, “Towards a ‘canonical’ agranular cortical microcircuit”, Frontiers in Neuroanatomy, vol. 8, ppp. 165, 2014.

[21] S. Geyer, M. Weiss, K. Reimann, G. Lohmann, and R. Turner, “Microstructural Parcellation of the Human Cerebral Cortex – From Brodmann’s Post-Mortem Map to in vivo Mapping with High-Field Magnetic Resonance Imaging”, Frontiers in Human Neuroscience, vol. 5, article 19, 2011.

[22] S. Clare, and H. Bridge, “Methodological issues relating to in vivo cortical myelography using MRI”, Human Brain Mapping, vol. 26, pp. 240–250, 2005.

[23] R. Turner, A. M. Oros-Peusquens, S. Romanzetti, K. Zilles, and N. J. Shah, “Optimised in vivo visualization of cortical structures in the human brain at 3 T using IR-TSE”, Magnetic Resonance Imaging, vol. 26, pp. 935–942, 2008.

[24] J. Dinse, N. Hartwich, M. D. Waehnert, C. L. Tardif, A. Schafer, S. Geyer, B. Preim, R. Turner, and P. L. Bazin, “A cytoarchitecture-driven myelin model reveals area-specific signatures in human primary and secondary areas using ultra-high resolution in-vivo brain MRI”, Neuroimage, vol. 114, pp. 71–87, 2015.

[25] A. Fracasso, S. J. van Veluw, F. Visser, P. R. Luijten, W. Spliet, J. J. M. Zwanenburg, S. O. Dumoulin, and N. Petridou, “Lines of Baillarger in vivo and ex vivo: Myelin contrast across lamina at 7T MRI and histology”, Neuroimage, vol. 133, pp. 163–175, 2016.

[26] M. D. Waehnert, J. Dinse, A. Schafer, S. Geyer, P. L. Bazin, R. Turner, & C. L. Tardif, “A subject-specific framework for in vivo myeloarchitectonic analysis using high resolution quantitative MRI”, Neuroimage, vol. 125, pp. 94–107, 2016.

[27] J. Annese, A. Pitiot, I. D. Dinov, and A. W. Toga, “A myelo-architectonic method for the structural classification of cortical areas”, Neuroimage, vol. 21, pp. 15–26, 2004.

[28] M. D. Waehnert, J. Dinse, M. Weiss, M. N. Streicher, P. Waehnert, S. Geyer, R. Turner, and P. L. Bazin, “Anatomically motivated modeling of cortical laminae”, Neuroimage, vol. 93, pp. 210–220, 2014.

[29] L. Tassi, N. Colombo, R. Garbelli, S. Francione, G. Lo Russo, R. Mai, F. Cardinale, M. Cossu, A. Ferrario, C. Galli, M. Bramerio, A. Citterio, and R. Spreafico, “Focal cortical dysplasia: neuropathological subtypes, EEG, neuroimaging and surgical outcome”, Brain, vol. 125, pp. 1719–1732, 2002.

